# Neocortical and hippocampal theta oscillations track audiovisual integration and replay of speech memories

**DOI:** 10.1101/2024.09.13.612424

**Authors:** Emmanuel Biau, Danying Wang, Hyojin Park, Ole Jensen, Simon Hanslmayr

## Abstract

“*Are you talkin’ to me?!*” If you ever watched the masterpiece “Taxi driver” directed by Martin Scorsese, you certainly recall the famous monologue during which Travis Bickle rehearses an imaginary confrontation in front of a mirror. While remembering this scene, you recollect a myriad of speech features across visual and auditory senses with a smooth sensation of unified memory. The aim of this study was to investigate how brain oscillations integrate the fine-grained synchrony between coinciding visual and auditory features when forming multisensory speech memories. We developed a memory task presenting participants with short synchronous or asynchronous movie clips focusing on the face of speakers engaged in real interviews. In the synchronous condition, the natural alignment between visual and auditory onsets was kept intact. In the asynchronous condition, auditory onsets were delayed to present lip movements and speech sounds in antiphase specifically with respect to the theta oscillation synchronising them in the original movie. We recorded magnetoencephalographic (MEG) activity to investigate brain oscillations in response to audiovisual asynchrony in the theta band. Our results first showed that theta oscillations in the neocortex and hippocampus were modulated by the level of synchrony between lip movements and syllables during audiovisual speech perception. Second, the accuracy of subsequent theta oscillation reinstatement during memory recollection was decreased when lip movements and the auditory envelope were encoded in asynchrony during speech perception. We demonstrate that neural theta oscillations in the neocortex and the hippocampus integrated lip movements and syllables during natural speech. We conclude that neural theta oscillations play a pivotal role in both aspects of audiovisual speech memories, i.e., encoding and retrieval.

## INTRODUCTION

The relative synchrony between visual and auditory inputs on theta rhythms during encoding has been shown to promote successful non-verbal audiovisual associations and performances in memory tasks (Clouter et al., 2017; Wang et al. 2018). In the brain, similar rhythmic patterns called theta oscillations originate from neuron assemblies and are thought to coordinate neurons’ firing in regions critical for episodic memory such as the hippocampus during perception (Buzsaki, 2010; Fries, Neuenschwader & Engel, 2001; Buzsaki, 2006; Buzsaki, 2013; Hanslmayr, Axmacher & Inman, 2019). Deep in the medial-temporal lobe, the hippocampus receives inputs from multiple sensory neocortices (Mayes, Montaldi & Migo, 2007; Moscovitch, 2008), and theta oscillations in this structure are thought to regulate at least two fundamental synaptic plasticity mechanisms supporting multisensory associations in episodic memory: In essence, the theta phase at which sensory inputs from the neocortex reach the ongoing theta oscillations in the hippocampus will determine long-term potentiation (LTP) or depression (LTD; Huerta and Lisman, 1995; Hölscher, Anwyl & Rowan, 1997). Further, the relative co-incident spiking occurring within short time intervals across multiple inputs leads to LTP/LTD via spike-timing dependent plasticity, i.e., optimal delays between pre- and post- synaptic spikes strengthen associations (Bi & Poo, 1998; Fell & Axmacher, 2011; Wang et al., 2023; Parish, Hanslmayr & Bowman, 2018). Based on this, we hypothesised that the role of audiovisual synchrony upon theta rhythms extends to speech memories as well. First, comparable rhythms imposed by the regular syllable onsets exist in continuous speech, which oscillate at a preferred theta rate (Giraud & Poeppel, 2012). In turn, previous EEG studies showed that the primary auditory cortex tracks and synchronises with speech envelope’s theta phase during perception (Peelle & Davis, 2012; Etard & Reichenbach, 2019; Gross et al., 2019; Park et al., 2015). Second, speech is often multisensory and the visual modality carries equivalent syllable information occurring at theta rate with the speaker’s lip movements (Biau et al., 2021; Ding et al., 2016; Bourguignon et al., 2019; Park et al., 2016). Unsurprisingly, the coherence between the phases of lip movements and auditory envelope during spontaneous audiovisual speech has been shown to be maximal in the theta band with a peak at ∼6 Hz corresponding to the syllable rate (Park et al., 2016; Chandrasekaran et al., 2009). Similarly, the visual cortex also tracks and synchronises with theta oscillations produced by the speaker’s lips during multisensory speech perception (Biau et al., 2021; Park et al., 2016). Consequently, during audiovisual speech perception listeners contemporarily process two streams of information which synchronise upon theta oscillations. This begs the question of whether the brain uses its internal theta oscillations to form integrated memories of audiovisual speech episodes. To address this question, our paradigm specifically manipulated the temporal alignment between visual and auditory information with respect to the dominant theta oscillation of each movie clip determined individually with mutual information analysis (**Figure 1A and 1B**). We predicted that (1) the specific asynchrony of the theta frequency aligning audiovisual information in the movies should decrease memory performance for multimodal speech associations. (2) Regions that integrate audiovisual information in the neocortex and hippocampus should track audiovisual synchrony with higher theta power for synchronous and lower theta power for asynchronous stimulation. Finally, we hypothesized that (3) upon retrieval speech signals are temporally replayed, in particular for stimuli which were synchronized during encoding.

**Figure 1.**
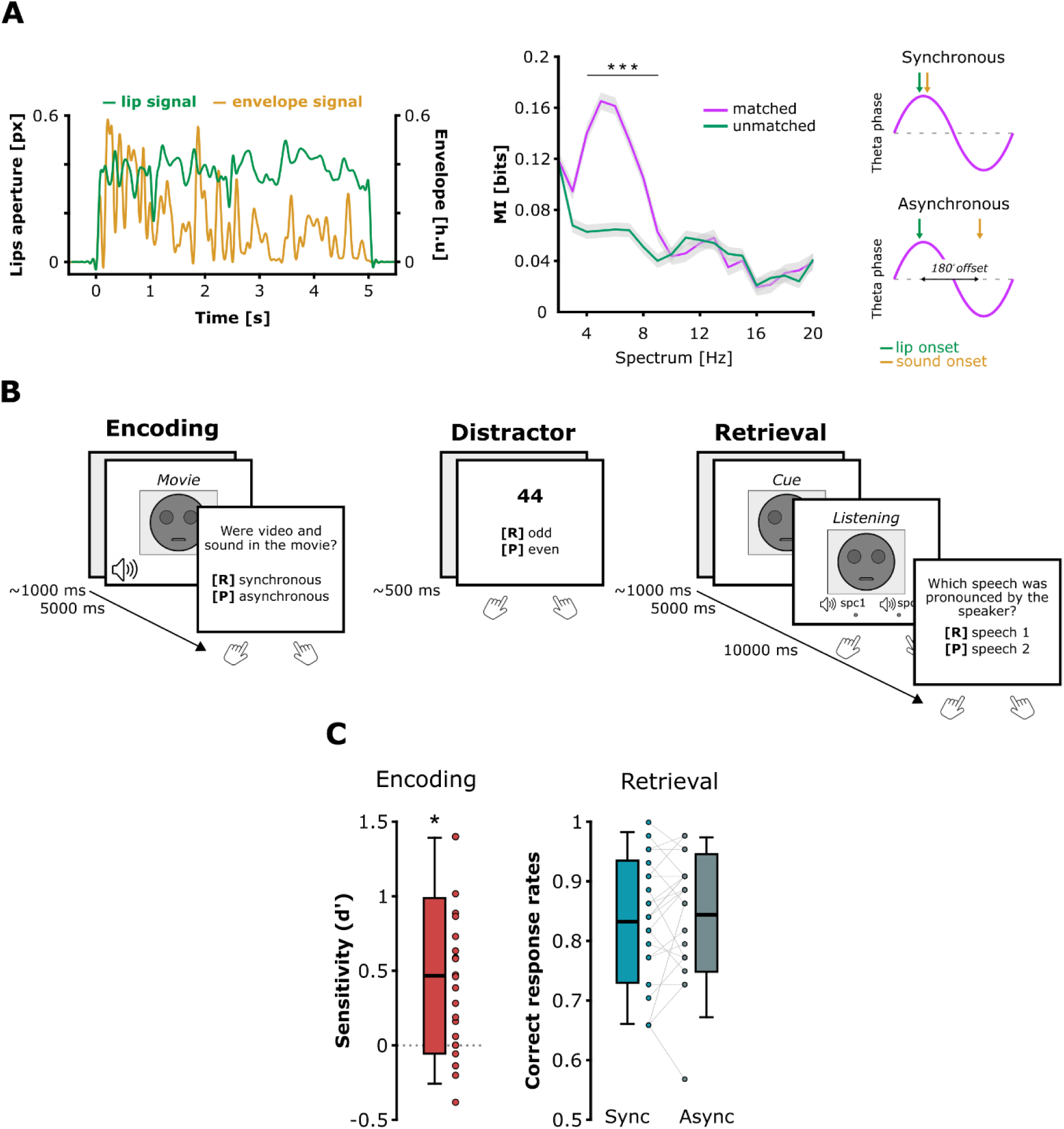
Movie stimuli, memory task and behavioural performances. (**A**) Left: Example of normalised lip movement (green line; pixel units) and auditory speech envelope (orange line; Hilbert units) time-series from the same movie. Centre: Coupling between the lip and auditory envelope signals in the movies (Mutual Information MI). The lip movements and speech envelopes predominantly synchronised on 4-8 Hz theta rhythms in the movies (purple line: mean ± standard error mean). This theta coupling disappeared when lip and speech envelope signals came from different movies (green line: mean ± standard error mean). Right: Illustration of the absolute theta timing manipulation induced between lip movement and auditory envelope onsets in order to create the synchronous and asynchronous conditions. (**B**) Example of one block of the procedure consisting in a sequence composed of encoding (speech movie presentations), distractor (rehearsal prevention) and retrieval (memory test on the movies perceived during the encoding). Participants saw a clear image of the face, which is drawn here for anonymity purpose. (**C**) Behavioural performance during encoding and retrieval. The sensitivity to theta audiovisual synchrony during movie encoding was assessed with the *d’*. Memory was assessed using the ratio of correct responses (i.e. where participants selected the correct speech stimulus pronounced by the speaker depicted during the cueing; 1 = 100% correct). The boxplots represent the mean ± standard deviation and the error bars indicate the 5th and 95th percentiles. The dots represent individual means. Significant contrasts are highlighted with asterisks (*p < 0.05; ***p < 0.001).

## RESULTS

### Audiovisual theta synchrony during speech encoding did not affect subsequent memory accuracy

During the encoding, participants memorised short audiovisual speech clips of speakers facing the camera. After each movie, they indicated with the keyboard whether lip movements and speech sounds were synchronous or asynchronous during the presentation. After a short distractor task, the participants completed the retrieval task. For each trial, they were cued with a picture depicting a previous speaker, and were instructed to silently remember the corresponding movie and its associated speech information. At the end of the visual cueing, the participants listened to two auditory speech stimuli and were instructed to indicate the auditory stimulus corresponding to the speaker cue (**Figure 1B** and Methods section). We computed the performance scores of the participants during the encoding and retrieval phases to address our first prediction (**Figure 1C**). In the memory task, participants correctly recalled the auditory speech associated with the speaker’s face during the retrieval (Synchronous: mean hit rate = 0.833 ± 0.103; Asynchronous: mean hit rate = 0.848 ± 0.099) and above chance level (i.e., 0.5 accuracy) in both synchronous (*t*(22) = 15.551; *p* < 0.001; Cohen’s *d* = 3.243; one-tailed) and asynchronous conditions (*t*(22) = 16.781; *p* < 0.001; Cohen’s *d* = 3.499; one-tailed). Nevertheless, results revealed no significant difference of accuracy between the two conditions behaviourally (*t*(22) = −1.005; *p* = 0.326; Cohen’s *d* = − 0.209; two-tailed). Therefore, the theta asynchrony between lip and sounds during speech encoding did not significantly impair the formation of speech memory (**Figure 1C right**). Participants’ sensitivity to theta timing during movie encoding was assessed with the *d’* from the audiovisual synchrony detection task (**Figure 1C left**). The averaged *d’* was close to zero (mean *d’* = 0.467 ± 0.522) but significantly positive (*t*(22) = 4.29; *p* < 0.001; Cohen’s *d* = 0.895; one-tailed). This result suggested that participants were able to distinguish between synchronous and asynchronous movies, although being far from perfect in doing that. Further analyses performed on the probability to respond “synchronous” in both conditions and the negative averaged bias *c* criterion corroborate this interpretation (supplemental information Figure S1).

### Neocortical and hippocampal oscillations integrate visual lip and sound information via theta phase

To test our second prediction, we first investigated how theta oscillations in the neocortex integrate cross-modal synchrony on the main frequency aligning visual and auditory speech features during movie perception. We contrasted theta power responses between the synchronous and asynchronous conditions at the whole brain level. Crucially, the power spectrum in every trial was first realigned on the frequency ±3 Hz corresponding to the peak of mutual information (MI) between the lip and auditory envelope signals determined in the movie analyses (**Figure 1A**). This step was critical in order to average all the trials together considering the main theta activity carried in each individual movie, before performing oscillation analyses (see Methods). The cluster-based analysis revealed a significant cluster which established that the perception of asynchronous audiovisual speech induced a decrease of theta power as compared to synchronous speech (*p* = 0.047, cluster size = 668.76, mean t-statistic within cluster = 2.164; Cohen’s *d_z_* = 0.451). The difference of theta power was localised bilaterally in the temporal and medial-temporal regions, in the right superior and inferior parietal lobules, and in the left frontal gyrus (**Figure 2A)**. These results established that the multisensory language network integrated the natural synchrony between lip movements and auditory envelope at theta oscillations (see **Figure S2** for theta power analysis during the encoding of synchronous and asynchronous movies separately). Here, we measured multisensory speech integration with theta power as it represents direct readout of the phase synchrony between the two sensory regions, which project into the associative neocortical regions (see **Figure S5 for phase coupling analysis**).

**Figure 2.**
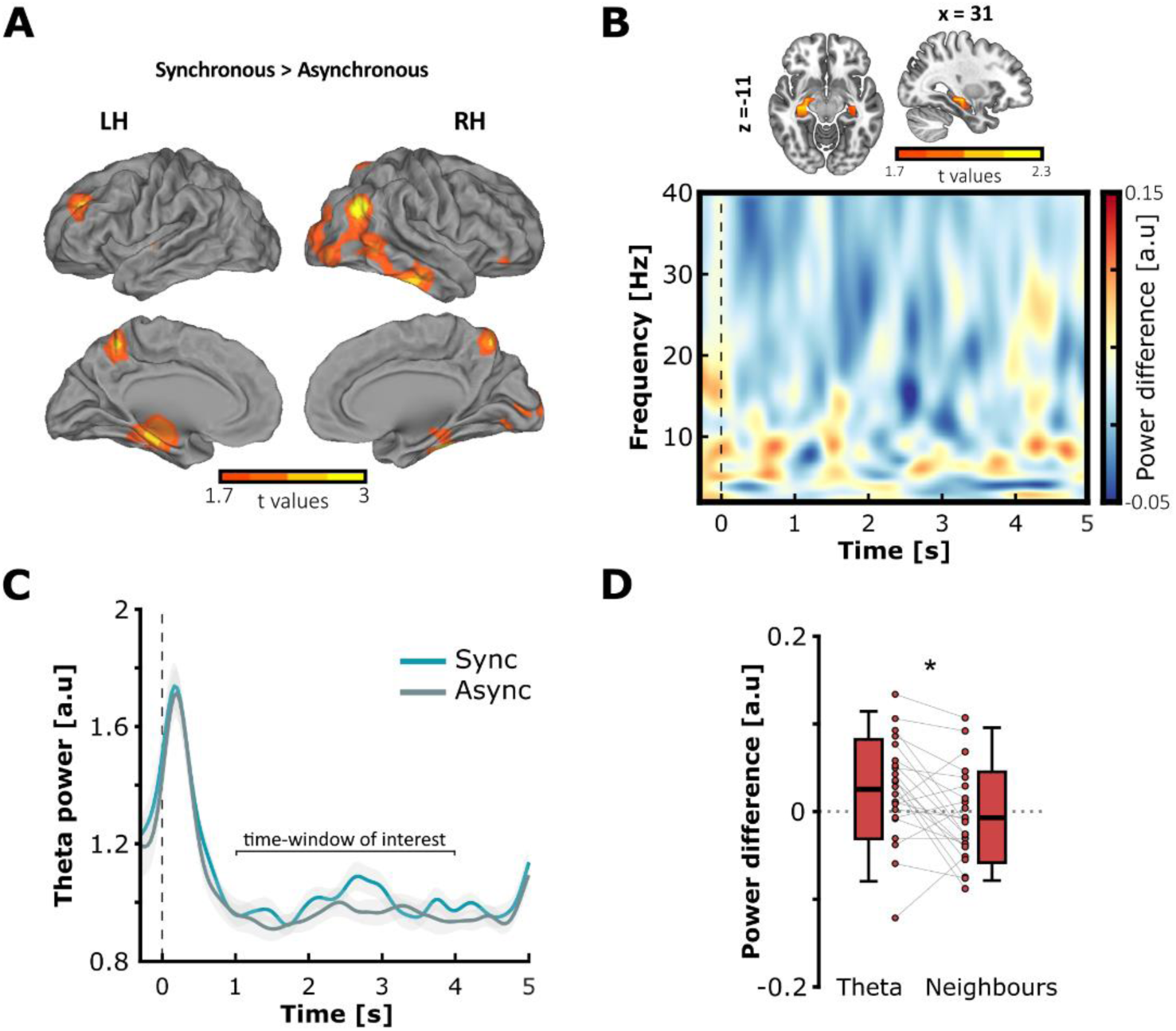
Brain oscillations integrate audiovisual theta timing in the neocortex and the hippocampus during movie encoding. (**A**) Source localisation of the difference of theta power in response to the audiovisual asynchrony in the movies (synchronous minus asynchronous). The perception of asynchronous movies induced a decrease of theta power as compared to synchronous movies, in cortical regions associated with language network (threshold at significant t values; cluster-corrected at α threshold = 0.05). (**B**) Difference of theta power in the hippocampus in response to the audiovisual asynchrony (*x* = 31; *y* = −19; *z* = −11; top) and time-frequency representation of the difference of spectral power in the hippocampus (bottom). Audiovisual asynchrony predominantly attenuated the power in the 4-8 Hz theta frequencies where lip movements and speech envelope were aligned in the synchronous condition during encoding. (**C**) Temporal modulation of theta power in the hippocampus during the encoding of synchronous (blue line: mean ± standard error mean) and asynchronous movies (grey line: mean ± standard error mean). (**D**) Mean difference of hippocampal power in the theta range versus neighbouring frequencies in the central time-window of interest. Theta oscillations in the hippocampus specifically integrated the audiovisual synchrony at the dominant frequencies (i.e. where lip movements and auditory envelopes were aligned in the movies). The boxplots represent the mean ± standard deviation and the errors bars indicate 5th and 95th percentiles. The dots represent individual means (* p<0.05).

Second, we tested whether theta oscillations integrated audiovisual synchrony in the hippocampus. The results shown in **Figure 2B (bottom panel)** suggest that audiovisual asynchrony indeed decreased the oscillatory responses predominantly in the low frequencies, overlapping with the theta frequency range. We then computed the mean difference of theta power between synchronous and asynchronous conditions, averaged within the central time-window during movie encoding (from +1 s to +4 s with respect to the onset) and across the virtual sensors from the hippocampus (left + right; **Figure 2B upper panel**). **Figure 2C** depicts the temporal modulation of theta oscillations, after realigning the frequency spectrum of the MEG in every single trial to the peak frequency in the physical stimulus (i.e. where lip movements and auditory envelope information showed maximal mutual information). Encoding of asynchronous movies induced a decrease of theta power compared to synchronous movies following the onset of the stimulus. The strong theta power increase at the start of the trial was likely induced by event-related-responses at the onset of the movies. It is worth noting that spontaneous hippocampal theta activity preceded the movie onsets, which is not surprising as theta oscillations dominate in the hippocampus (Buzsaki, 2006; Griffiths et al., 2019). Most relevant to our hypothesis, we found that hippocampal theta power was modulated by the synchrony/asynchrony of the movies (mean theta power difference = 0.025 ± 0.057; *t*(22) = 2.11; *p* = 0.023; Cohen’s *d* = 0.44), confirming that theta oscillations in the hippocampus temporally integrate visual and auditory information during speech encoding. To verify whether hippocampal theta oscillations tracked specifically the dominant activity aligning visual and auditory information in the movies, we compared the magnitude of hippocampal theta power difference between the 4-8Hz range of interest and neighbouring frequency bands (**Figure 2D**). To do so, each original theta frequency was divided and multiplied by two times the golden mean (1.618) to determine the same number of frequencies with no shared harmonic in the low and high neighbouring bands surrounding the theta range (Pletzer, Kerschbaum and Klimesch, 2010; see Method section). In contrast with the theta band, a one-sample t-test showed that the difference of mean power in the neighbouring frequencies was not significantly different from zero (mean power difference = −0.007 ± 0.052; *t*(22) = −0.674; *p* = 0.746; Cohen’s *d* = 0.14). A paired-sample t-test confirmed that the theta power difference was significantly greater in the 4-8Hz frequencies of interest than in the neighbouring frequencies (*t*(22) = 2.866; *p* = 0.005; Cohen’s *d* = 0.587). Altogether, the results show that theta oscillations integrate the fine-grained syllabic organisation of audiovisual speech during perception. Further, this organisation was maintained downstream in the hippocampus, which may promote the formation of audiovisual associations in memory.

### Audiovisual theta synchrony during encoding decreased the accuracy of speech replay from memory

Finally, we examined whether theta patterns from auditory cortices reflect the accuracy of replay of the speech envelope from memory, i.e. when participants recalled auditory information cued by the face of the speaker (**Figure 3**). To this end, we first needed to confirm that neural theta oscillations tracked the dominant activity conveyed in the auditory speech envelope during encoding, i.e. initial movie perception. To this end, we expected to find a theta ‘tracking’ (i.e. alignment of theta phase with physical stimulus) in the primary auditory cortex (Heschl’s gyrus) during encoding.

**Figure 3.**
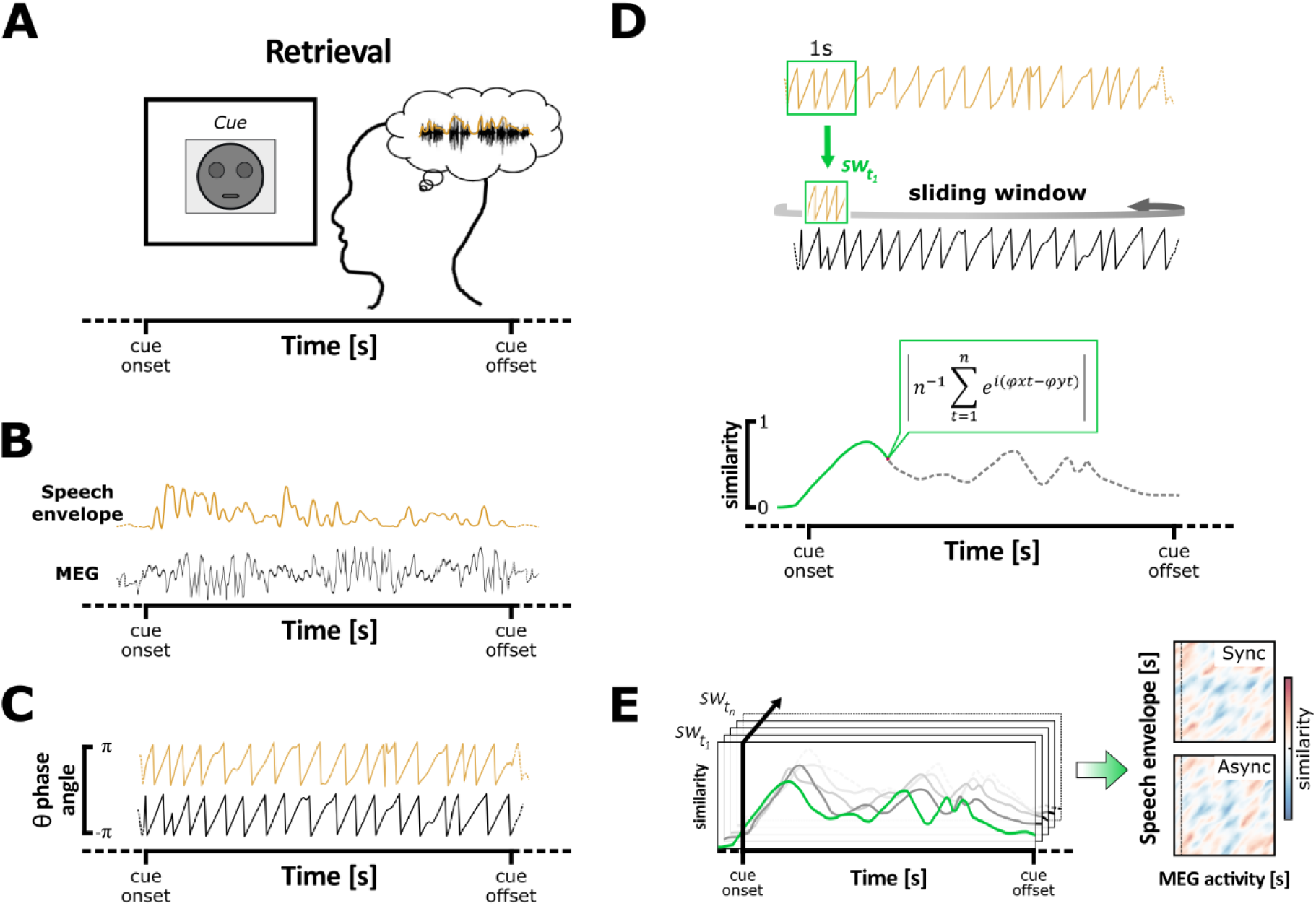
Measure of speech replay accuracy during retrieval. (**A**) During successful retrieval, participants recalled the auditory features associated with the speaker’s face, theoretically leading to mentally replaying *verbatim* the speech memorised during movie encoding. (**B**) In this case, the brain activity in the auditory cortex (MEG; black line) reinstates the theta oscillatory patterns conveyed by the auditory speech envelope of the movie (Speech envelope; orange line). (**C**) The accuracy of speech reinstatement is measured with the level of phase similarity between the theta oscillation of the stimulus auditory envelope, and the neural theta oscillation produced replay of speech memory. To do that, the phase of the dominant theta frequency was extracted from the auditory envelope of the stimulus (orange line) and the MEG signal at the auditory sensors (black line). (**D**) The phase similarity was estimated between the first sliding window (sw) of one second containing the theta phase of the auditory envelope centred on its first time-point (sw*_t1_*), and every time-point of the MEG signal (top). The phase similarity was represented with a single value ranging between 0 and 1 at each time-point of the MEG signal (bottom). (**E**) This operation was repeated by shifting the sliding window to the next time-point (sw*_t2_*) of the auditory envelope signal and so on until its last time-point (sw*_tn_*), in order to compute the phase similarity between the two signal dimensions over time (left). Theta phase similarity was then averaged across windows, trials and sensors of interest for statistical analyses (right). Adapted from Michelmann et al. (2016).

Figure 4A depicts the statistical difference of mean theta phase similarity averaged across pairs of corresponding MEG and envelope signals from the same trials (i.e., matched) minus the mean theta phase similarity averaged across random pairing of signals coming from different trials (i.e., unmatched) during the encoding of synchronous and asynchronous movies. The results revealed a significant cluster when testing the phase similarity difference between matched and unmatched signals in both the synchronous (*p* = 0.003, cluster size = 9.321x10^3^, mean t-statistic within cluster = 2.936; Cohen’s *d_z_* = 0.612) and asynchronous condition (*p* < 0.001, cluster size = 1.402x10^4^, mean t-statistic within cluster = 4.259; Cohen’s *d_z_* = 0.888). These results showed that during the encoding of audiovisual movies, neural theta oscillations tracked the dominant activity conveyed by the auditory speech signal across a broad neocortical network. Although not exclusive, the greatest theta tracking was situated bilaterally in the temporal regions including the primary auditory cortex (left and right Heschl’s gyrus), as well as pre- and post-central areas. This pattern appeared to be clearer during the encoding of audiovisual synchronous movies as compared to asynchronous ones, suggesting that the asynchrony between the lip movements and the speech envelope indeed affected the natural tracking of the auditory signal during encoding (the equivalent analysis performed between the MEG activity and the lip movement signals are reported in **Figure S4,** which was not our principal aim as participants were instructed to recall auditory information during the visual cueing). Next, we aimed to establish whether neural theta oscillations effectively indexed the reinstatement of the dominant theta dynamics carried in the auditory envelope during retrieval, independently from the condition of encoding (i.e., if they provide a valid signature of memory replay). We computed the theta phase similarity between every time-point of the auditory cortex (left + right) and corresponding speech envelope signals from the synchronous and asynchronous conditions together, which was tested against permuted data. The cluster-based analysis revealed that phase similarity was significantly greater than chance-level, establishing that neural theta phase reflects replay of auditory speech envelopes of encoded movies (*p* < 0.001, cluster size = 7.554x10^4^, mean t-statistic within cluster = 4.388; Cohen’s *d_z_* = 0.915; Figure 4B). This effect was dominated by enhanced replay at the beginning and end of speech envelopes, suggesting that these parts of the signal were most informative for participants during movie encoding. To control for any purely onset-/offset-driven effect of the stimuli, our analysis contrasted the phase similarity averaged across pairs of corresponding MEG and speech envelope signals against permuted data generated by randomly pairing each MEG signal with an envelope signal selected from a different trial. Therefore, any effect purely driven by the onset or offset would have been preserved in both the corresponding and permuted data, which would have cancelled out any early or late difference reported in Figure 4B. On the other hand, the observed patterns of replay possibly reflect some primacy- and recency-driven phenomenon, illustrating that participants better encoded, and therefore replayed predominantly the beginning and the end of stimuli (Glanzer & Cunitz, 1966).

**Figure 4.**
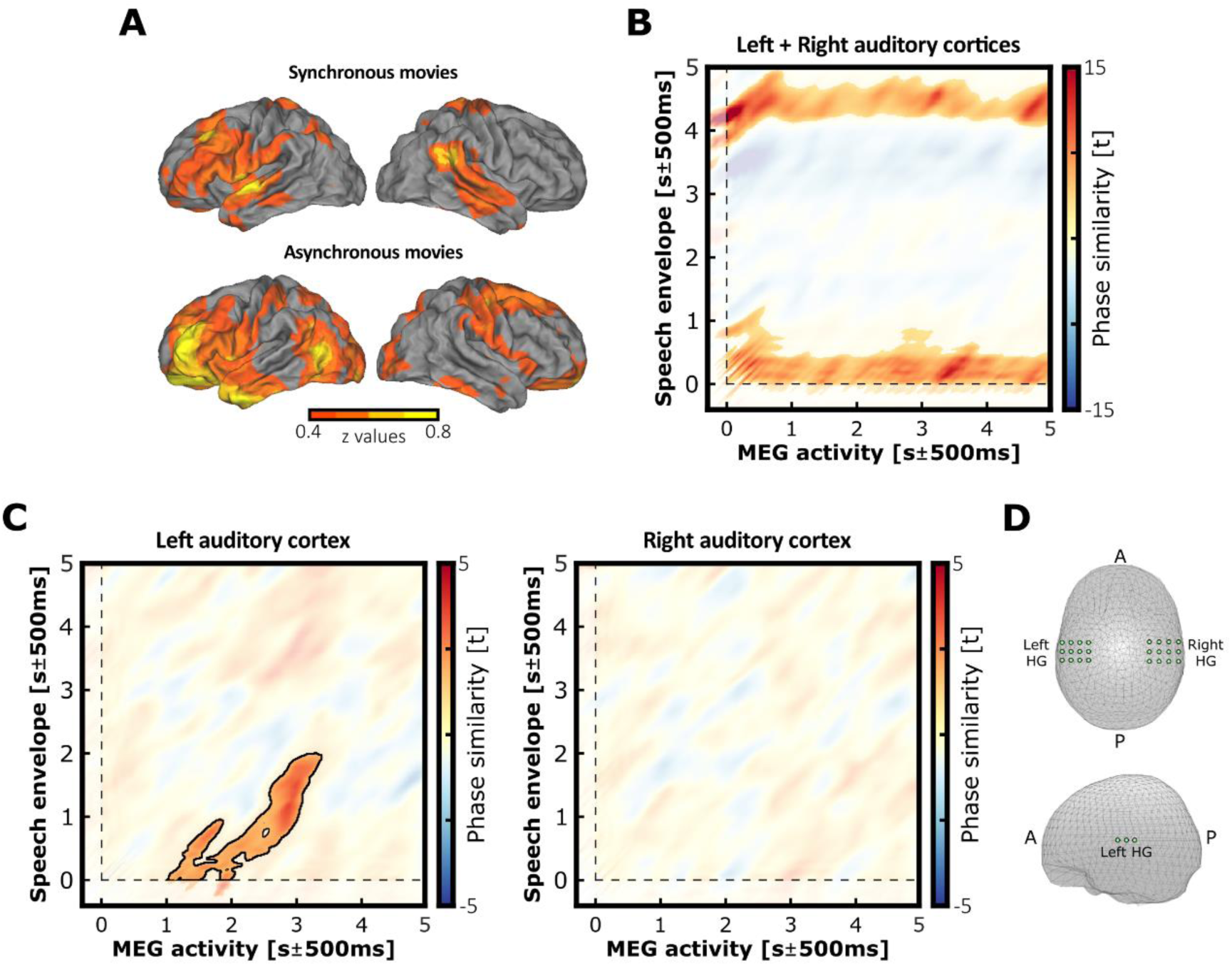
Theta oscillations track and reinstate the dominant activity carried in the auditory envelope. (**A**) Source localisation of theta phase similarity between the brain oscillations and auditory speech envelope during synchronous or asynchronous movie encoding (threshold at significant t values and normalized for visualisation). The tracking of the dominant theta oscillations carried by the auditory speech information appeared more consistent bilaterally at the expected auditory regions when lip movements and speech sounds were perceived in synchrony compared to asynchrony. (**B**) Theta phase similarity between the theta oscillations from auditory cortices (left + right; X-axis) and the theta oscillations conveyed by auditory speech envelope (Y-axis) during retrieval. Y-axis represents time for the speech envelope, x-axis represents time for the MEG theta phase. Theta oscillations in the auditory cortex reflect the reinstatement of auditory speech envelopes when participants successfully recalled auditory information. The time-points from the significant cluster are depicted in full colour, while non-significant data are masked with transparency. (**C**) Theta phase similarity difference between the synchronous versus asynchronous conditions in the left and right auditory cortices during successful retrieval. The accuracy of the reinstatement was modulated by the audiovisual asynchrony in the left but not the right auditory cortex. The time-points from the significant cluster are depicted in full colour and outlined, while non-significant data are masked with transparency. (**D**) Virtual sources localised in the left and right auditory cortices used for the phase similarity analysis (respectively the left and right Heschl’s gyrus; LHG and RHG).

Lastly, we compared whether there was a difference in the accuracy of replay of auditory speech envelopes between synchronous and asynchronous movies in auditory cortices. To this end, the theta phase similarity between every time-point of the MEG and corresponding speech envelope signals was computed in the left and right auditory cortices during the successful retrieval (Figure 4C). Results revealed higher theta phase similarity for synchronous compared to asynchronous movies in the left auditory cortex (*p* = 0.023, cluster size = 1.22x10^4^, mean t-statistic within cluster = 2.226; Cohen’s *d_z_* = 0.464). This result confirmed that the accuracy of speech replay in the left auditory cortex was decreased when movies were encoded in the asynchronous as compared to the synchronous condition (Figure 4C **left**). In contrast, the cluster-based permutation test revealed no significant difference between the synchronous and asynchronous conditions in the right auditory cortex (Figure 4C **right**). For completeness, the difference of theta phase similarity between conditions was further tested at the whole brain level within a 1000-ms widow centred on the maximum t-value coordinate from the above significant cluster (respectively 2.891 s ± 500 ms in the MEG dimension and 1.234 s ± 500 ms in the envelope dimension with respect to signal onsets). Nevertheless, results revealed no significant cluster in both auditory cortices (**Figure S6**). Additional analyses of hippocampal theta power during retrieval are reported in the supplemental information (**Figure S3**). Altogether, our results established that the auditory cortex reinstated theta phase patterns imposed by speech in the dominant frequency in the previous movies. This suggests that the theta dynamics were encoded as relevant speech features contributing to memory replay.

## DISCUSSION

In everyday life, we form an abundant amount of speech memories during casual conversations or when we see a movie. Replaying later what someone said and how they said it without much effort requires the brain to form an integrated representation of multisensory speech in memory. Theta oscillations in the neocortex and the hippocampus have shown to play an important role for episodic memory formation. Moreover, syllable information organises speech on equivalent theta oscillations that align lip movements and auditory envelope. We go beyond previous work in showing that theta synchrony between speech and neuronal activation in neocortex and hippocampus reflects whether audiovisual speech was encoded in synchrony or not. Further, we found that audiovisual asynchrony during encoding affects the accuracy at which speech stimuli are later replayed from memory.

Decreasing audio-visual theta phase alignment in the movies attenuated neural responses in right temporal regions including posterior middle temporal and superior temporal gyri, which are commonly associated with multisensory integration (Calvert, Campbell & Brammer, 2000; Callan et al., 2004; Macaluso et al., 2004; Meyer et al., 2004; Campbell, 2008; Nath and Beauchamp, 2012; Jansma et al., 2014). These areas often exhibit early superadditive activations within the first 160 ms in response to congruent audiovisual speech perception compared to unimodal speech (Gao et al., 2023; Salmi et al., 2017; Besle et al. (2008); Besle et al., 2009; Molholm et al., 2002). Further studies also reported that audiovisual integration takes place in both auditory and visual areas of the neocortex (Ghazanfar & Schroeder, 2006; Petro, Paton & Muckli, 2017; Raij et al., 2010). For instance, Park et al. (2018) demonstrated that 3-7Hz theta oscillations in the posterior superior temporal gyrus/sulcus (STG/S) encoded common features carried in auditory and visual inputs during audiovisual speech perception. Lip and sound onsets were misaligned by up to 125 ms in the stimuli, and the decrease of specific theta power found in the right temporal and visual regions suggests that theta asynchrony reduced audiovisual integration in both sensory and associative areas. Therefore, our results confirmed that the neocortex integrates lip-sound temporal mapping upon theta rhythm. Finally, the STG has been shown to encode acoustic-phonemic features during speech processing (Yi, Leonard & Chang, 2019; Giraud & Poeppel, 2012). Here, it is possible that desynchronising lips and sounds affected acoustic-phonemic feature processing as well, because their dynamics depend on the syllable timing. This idea is corroborated by the theta activity tracking of the auditory envelope during movie encoding, which was found in the bilateral STG and was attenuated in the right hemisphere during the encoding of asynchronous movies. It is also possible that the audiovisual theta asynchrony imposed in the movies added a “visual noise” that shaped speech sound processing in the auditory associative STG/S regions (Gwilliams et al., 2018; Holdgraf et al., 2016).

Next, we showed that the hippocampus integrates the natural theta timing in speech as well. This was reflected by a decrease of power specifically in those theta frequencies where multisensory syllable information was asynchronous during movie encoding. Previous studies demonstrated that encoding of asynchronously flickering video-sound associations at theta predicted worse memories compared to synchronously flickering associations (Clouter et al. 2017; Wang et al., 2018). From the delay observed between the theta oscillation across cortical sensory areas, these studies hypothesized that the same theta timing was maintained downstream in the hippocampus to synchronise the responding neural activities and to promote audiovisual memory associations. Using MEG, the present study demonstrates hippocampal theta power modulations in response to the timing between multisensory speech features. This result was anticipated because during the encoding, the hippocampus receives multisensory information from the neocortex via the perforant pathway of the entorhinal region (Suzuki, 1996; Mayes, Montaldi & Migo, 2007; Moscovitch, 2008). In rodents and non-human primates, the auditory and visual sensory as well as associative cortical areas project to the parahippocampal and perirhinal cortices, which send information inputs to the entorhinal cortex (Witter, 1989; Witter & Amaral, 1991) which passes it on to the hippocampus. Cancelling theta synchrony in the frequency organising audiovisual speech movies is likely to affect multisensory integration normally taking place in neocortical associative areas, which would then pass on the misaligned information downstream to the hippocampus. It is worth noting that although MEG has traditionally been challenged with detecting activity originating in deep sources, recent studies have now established that MEG is indeed sensitive to signals from the hippocampus (Joensen et al., 2023; Alberto et al., 2021; Ruzich et al., 2019).

The results show that the neocortical theta phase tracked theta phase information carried in the speech envelope during movie encoding. Strikingly, this theta phase activity is later reinstated in the auditory cortices during subsequent retrieval. Interestingly, the audiovisual synchrony at encoding was associated with higher accuracy of speech replay at retrieval in the left auditory cortex. Using a similar analytic approach, a previous study showed the replay of auditory phase patterns at 8 Hz of music stimuli (Michelmann, Bowmann & Hanslmayr, 2016). In a verbal memory task, Yaffe et al. (2014) reported that theta oscillations from the bilateral temporal lobes mediated neural reinstatement associated with successful paired-word retrieval. However, words were encoded in isolation and theta activity did not encode dynamic features of the event *per se*. Here, we extended these results with continuous speech stimuli by showing that the brain replays temporal patterns that organised speech information during encoding, and that this replay depends on the temporal synchrony at initial encoding. Theta oscillations are thought to provide the optimal timing to promote LTP during the formation of new multisensory associations (Pavlides et al., 1988; Huerta and Lisman, 1995; Holscher et al., 1997; Buzsaki, 2002; Hyman et al., 2003; Wang et al., 2023). The less accurate theta reinstatement observed when participants recalled asynchronous movies may reflect the weaker reactivation of memory traces as compared to synchronous movie memories. We speculate that misaligning lip movements and speech sounds induced a subtle desynchronization between responding downstream assemblies in the hippocampus, which affected their optimal theta timing and decreased the likelihood of LTP during encoding. Nevertheless, recall performance in the subsequent memory test was not decreased by the audiovisual asynchrony during movie encoding. Therefore, our results have not confirmed our first prediction yet.

If audiovisual asynchrony reduced the accuracy of replay of speech stimuli, then why did it not affect memory performance? We see two possible answers. First, participants were able to distinguish between synchronous and asynchronous movies during encoding (see Figure 1C). Therefore, it is possible that asynchronous movies attracted listeners’ attention and artificially “boosted” audiovisual associations. Second, the memory task required participants to simply choose between two speech stimuli, which may not be sensitive to measure the accuracy of memory replay. For instance, a participant could reach very high levels of performance if they simply remember the rough content of what the speaker said (i.e., the speaker said something about “home-works”) as opposed to a fine-grained temporal representation of the speech content. Future investigations should aim to improve the granularity of the memory task by testing participants on specific speech features which would be reflected in theta phase.

## CONCLUSION

We demonstrated that neural theta oscillations in the neocortex and the hippocampus integrated lip movements and syllables during natural speech. Further, the replay of the temporal features encoded in the speech movies were mediated by the same theta oscillations. The accuracy of theta reinstatement during memory recollection was affected by the synchrony between lip movements and the auditory envelope during speech encoding. We conclude that neural theta oscillations play a pivotal role in both aspects of audiovisual speech memories, i.e., encoding and retrieval.

## SUPPLEMENTAL INFORMATION

Supplemental information contains X figures, one movie clip, and can be found at XXX.

## AUTHOR CONTRIBUTION

E.B and S.H designed the experiments and paradigms. E.B and D.W collected the data. E.B analysed the data. E.B, D.W, H.P, O.J and S.H wrote the paper. All the authors discussed the results and commented on the manuscript.

## ACKNOWLEDGEMENT

This work was supported by a Sir Henry Wellcome Fellowship awarded to E.B. (Grant reference number: 210924/Z/18/Z), as well as grants from the ERC (Consolidator Grant 647954) and ESRC (ES/R010072/1) awarded to S.H.

## KEY RESOURCES TABLE

| REAGENT or RESOURCES | SOURCES | IDENTIFIER |
| --- | --- | --- |
| **Software and algorithms** |  |  |
| MATLAB | www.mathworks.com | R2018 |
| Psychophysics Toolbox | psychtoolbox.org | 3 |
| Fieldtrip | www.fieldtriptoolbox.org | 20171231 |
| Circular Statistics Toolbox | www.mathworks.com/matlabcentral/fileexchange/10676-circular-statisticstoolbox-directional-statistics | 1.21.0.0 |
| SPM | Wellcome Trust Centre for Neuroimaging | 8 |
| Caret | Washington University School of Medicine | 5.65 |
| MRICroGL | www.nitrc.org/projects/mricrogl | 1.2.20190902 |
| **Other** |  |  |
| TRIUX MEGIN | www.megin.com | MEG scanner |
| 3-Tesla Siemens PRISMA | www.siemens-healthineers.com | MRI scanner |
| ER-3C system | www.etymotic.com | MEG compatible earphones |
| Fastrack | www.polhemus.com | Electromagnetic digitizer |

## RESOURCE AVAILABILITY

### Lead contact

Further information and requests for resources and reagents should be directed to and will be fulfilled by the lead contact, Emmanuel Biau.

### Material availability

This study did not generate new unique reagents.

### Data and code availability

The data sets have been deposited at https://osf.io/4nh5w/ and will be made available after publication. All original codes have been deposited at https://osf.io/4nh5w/ and will be made available after publication. Any additional information related to the present article will be available from the lead contact upon request.

## EXPERIMENTAL MODEL AND SUBJECT DETAILS

Thirty healthy participants took part in the experiment (mean age = 24.17 years ± 2.71; 9 females). All participants were English native speakers and right-handed. All of them reported normal or corrected-to-normal vision and hearing. All participants received financial reimbursement for taking part in the experiment (£10 per hour). Three participants were excluded because of technical issues affecting the presentation of sounds during the experiment; four additional participants were excluded because of excessive noise in the data. This left twenty-three data sets for analysis. All participants gave written informed consent. Ethical approval was granted by the University of Birmingham Research Ethics Committee (ERN_18-0226AP10) complying with the Declaration of Helsinki.

## METHOD DETAILS

### Movie stimuli

Eighty-eight five-second audiovisual movies were extracted from natural face-to-face interviews freely accessible and downloaded from YouTube (www.youtube.com). Satisfying movies containing meaningful content (i.e., at least one complete sentence, speaker facing toward the camera) were edited using Shotcut (Meltytech, LLC). For each movie, the video and the sound were exported separately (Video: .mp4 format, 1280 x 720 resolution, 30 frames per second, 200 ms linear ramp fade in/out; Audio: .wav format, 44100 Hz sampling rate, mono). Edited stimuli are available on https://github.com/EmmanuelBiau/SilentLip_perception and were validated in a previous study (Biau et al., 2021).

### Lip movement detection

Lips contour signal was extracted for each video using in-house Matlab codes. We computed the area information (area contained within the lips contour), the major axis information (horizontal axis within lip contour) and minor axis information (vertical axis within lip contour) as described in Park et al. (2016). In the present study, we used vertical aperture information of the lips contour to establish the correspondence between lips and auditory speech at the syllabic rate (i.e. aperture between the superior and inferior lips). The lips time-series was resampled at 250 Hz for further analyses with corresponding auditory speech envelope (Figure 1A **left).**

### Auditory speech envelope

The amplitude envelope of each movie sound was computed using in-house Matlab codes (Park et al., 2016; Chandrasekaran et al., 2009). First, eight equidistant frequency bands spanning on the cochlear map in the range 100-10,000 Hz were constructed (Smith et al., 2002). Narrow band sound signals were then band-pass filtered with a fourth-order Butterworth filter (forward and reverse). Absolute Hilbert transform was applied to obtain amplitude envelopes for each narrow band. These signals were then averaged across bands and resulted in a unique wideband amplitude envelope per sound signal. Each final signal was resampled to 250 Hz for further theta correspondence analyses (Figure 1A **left).**

### Mutual information between lip movements and auditory speech envelope

To identify the dominant activity aligning visual and auditory syllable information, we determined at which theta frequency the auditory and visual speech signals showed significant dependencies for each stimulus. To do so, we examined the audiovisual speech frequency spectrum (1 to 20 Hz) and computed the mutual information (MI) between the minor axis information and speech envelope signals sampled at 250 Hz. MI measures the statistical dependence between two variables with no prior hypothesis, and with a meaningful effect size measured in bits (Ince et al., 2017; Shannon, 1948). We applied the Gaussian Copula Mutual Information (GCMI) approached described in Ince et al. (2017) in which the MI between two signals corresponds to the negative entropy of their joint copula transformed distribution. This method provides a robust, semiparametric lower bound estimator of MI by combining the statistical theory of copulas together with the closed-form solution for the entropy of Gaussian variables, allowing good estimation over circular variables, like phase as well as power. For each movie, the complex spectrum is normalized by its amplitude to obtain a 2D representation of the phase as points lying on the unit circle for both the lip movements and auditory envelope time-series. The real and imaginary parts of the normalized spectrums are rank normalized separately and the phase dependence for each frequency between the two 2D signals is estimated using the multivariate GCMI estimator giving a lower bound estimate of the MI between the phases of the two signals. Here, we applied the GCMI analyses to determine to select and validate the movie clips: (1) We computed MI between matching lip and auditory envelope signals (i.e., the lip movement and auditory envelope signals were computed from the same audiovisual speech clip). (2) For each audiovisual stimulus, we performed a peak detection on the MI spectrum (1-20 Hz) to determine which specific frequency aligned best lip movements and auditory envelope in every movie (mean frequency MI peak across stimuli: 5.750 ± 1.341 Hz; 20 videos with a peak of MI at 4 Hz; 22 videos with a peak of MI at 5 Hz; 16 videos with a peak of MI at 6 Hz; 20 videos with a peak of MI at 7 Hz; 10 videos with a peak of MI at 8 Hz). (3) We computed MI between non-matching signals for every stimulus (i.e., the lip movement signal of each stimulus was randomly paired with the auditory envelope of another stimulus before performing MI analysis). T-test comparisons of the averaged MI across the 1-20 Hz spectrum between matching vs. non-matching stimuli confirmed a significant difference only in the 4-9 Hz theta band (4hz: t(19) = 5.846; p = 8.580x10-7; 5hz: t(19) = 6.561; p = 3.672 x10-8; 6hz: t(19) = 5.724; p = 1.452x10-6; 7hz: t(19) = 4.810; p = 6.272x10-5; 8hz: t(19) = 5.064; p = 2.270x10-5; 9hz: t(19) = 2.976; p = 0.038. For the remaining frequencies of the spectrum (i.e., from 1 to 3 Hz and from 10 to 20 Hz), the MI was not different between matching vs. non-matching stimuli. All the p-values were Bonferroni-corrected for multiple comparisons. These results are well in line with previous studies using coherence or MI measures (Park et al., 2016; Chandrasekaran et al., 2009), confirming that lip movements and auditory speech signals aligned preferentially in the theta band as compared to the rest of the spectrum, which validates the selected audiovisual speech stimuli (Figure 1A **centre**).

### Audiovisual theta synchrony in movie stimuli

For the present study, we created two versions of audiovisual speech from the 88 video clips (Figure 1A **right**): (1) The synchronous version in which the natural synchrony between video and sound signals from the original clips was maintained intact. Therefore, lip movements and speech envelope fluctuations aligned on the preferred theta oscillation imposed by syllable onsets and determined individually for each stimulus (c.f., analysis described above). (2) To create the asynchronous version, we shifted the onset of the auditory signal with respect to the visual signal by 180° of the preferred theta oscillation (i.e. peak theta frequency of MI). Therefore, this delay (in absolute time) depended on the preferred theta oscillation aligning lip movements and auditory envelope signals determined in the previous MI analysis. For instance, if lip movements and speech envelope aligned at 4 Hz in the movie *n*, the onset of the auditory signal was delayed by 125 ms with respect to its original timing to induce an offset with the visual signal onset corresponding to 180° in the theta phase. Instead, if lip movements and speech envelope aligned best at 5 Hz in another movie *n+1*, the auditory signal onset was delayed by 100 ms to create the asynchronous version of this movie *n+1*. The exact same logic was applied to create the asynchronous version of other audiovisual clips in which lip movement and auditory envelope signals aligned best at 6,7 or 8 Hz, by inducing a delay of 83, 71 or 63 ms, respectively. Finally, the natural order of signal presentation was always kept across stimuli and conditions, i.e., lip aperture leading its corresponding sound.

### Experimental paradigm

Participants seated comfortably in the MEG gantry, approximatively 150 cm away from the projection screen. The task was presented on a middle-grey screen (1920x1200 pixel resolution) using Psychophysics Toolbox-3 (Brainard & Vision, 1997). The sound was presented between 70 and 80 dB through MEG-compatible insert earphones (Etymotic Research, Elk Grove Village, IL). The experimental paradigm consisted of repeated blocks, with each block being composed of three successive tasks: encoding, distractor and retrieval tasks (Figure 1B). During the encoding, participants were presented with a series of audiovisual speech movies and performed an audiovisual synchrony detection. Each trial started with a brief fixation-cross (jittered duration: 1000-1500 ms) followed by the presentation of a random synchronous or asynchronous audiovisual speech movie (5 s). After the movie end, participants had to determine whether video and sound were presented in synchrony or asynchrony in the movie, by pressing the index finger (“synchronous”) or the middle finger (“asynchronous”) button of the response device as fast and accurate as possible. The next trial started after the participant’s response. After the encoding, the participants did a short distractor task. Each trial started with a brief fixation-cross (jittered duration: 1000-1500 ms) followed by the presentation of a random number (from 1 to 99) displayed at the centre of the screen. Participants were instructed to determine as fast and accurate as possible whether this number was odd or even by pressing the index (“odd”) or the middle finger (“even”) button of the response device. Each distractor task contained twenty trials. The purpose of the distractor task was only to clear memory up. After the distractor task, the participants performed the retrieval task to assess their memory. Each trial started with a brief fixation-cross (jittered duration: 1000-1500 ms) followed by the presentation of a static frame depicting the face of a speaker from a movie attended in the previous encoding. During this visual cueing (5 s), participants were instructed to recall as accurately as possible every auditory information previously associated with the speaker’s speech during the movie presentation. At the end of the visual cueing, participants were provided the possibility to listen two auditory speech stimuli: one stimulus corresponded to the speaker’s auditory speech from the same movie (i.e., matching). The other auditory stimulus was taken from another random movie (i.e., unmatching). Participants chose to listen each stimulus sequentially by pressing the index finger (“Speech 1”) or the middle finger (“Speech 2”) button of the response device. The order of displaying was free. At the end of the second auditory stimulus, participants were instructed to determine as fast and accurate as possible which auditory speech stimulus corresponded to the speaker’s face frame, by pressing the index finger (“Speech 1”) or the middle finger (“Speech 2”) button of the response device. The next retrieval trial started after the participant’s response. After the last trial of the retrieval, participants took a short break, before starting a new block (encoding-distractor-retrieval).

### Staircase procedure and speech condition counterbalancing

In total, the paradigm presented 88 audiovisual speech stimuli with a staircase procedure adapting online to individual performances in order to limit any ceiling effect in the memory test: the encoding task of the first block always contained 16 trials. Then, the length of the encoding task in the next block depended on the accuracy in the previous retrieval task (i.e., the percentage of auditory speech stimuli correctly retrieved with the visual cue) and was determined as follows: when accuracy in the retrieval *n* was between 70 and 80 percent, the encoding *n+1* remained the same. If memory accuracy in the retrieval *n* was above 80 percent, the length of the encoding *n+1* was increased with 4 more trials to increase difficulty in retrieval *n+1*. If accuracy in the retrieval *n* was below 70 percent, the length of the encoding *n+1* was decreased with 2 less trials to decrease difficulty in retrieval *n+1*. During the encoding, half of the total audiovisual stimuli were presented in the synchronous condition (44 trials) and the other half was presented in the asynchronous condition (44 trials). To control for material effects, i.e. some stimuli might be more memorable than others, the assignment of stimuli to the synchronous or asynchronous condition was counterbalanced across participants (e.g., the synchronous version of the movie *n* was presented to participant *X* and the asynchronous one to participant *X+1*).

### Behavioural performances and analysis

(1) Audiovisual synchrony detection: trials for which participants correctly detected audiovisual synchrony between video and sound in the synchronous condition were labelled as “hit” (i.e., participants responded “synchronous” in the synchronous audiovisual speech movies). Trials for which participants wrongly responded synchronous in the asynchronous condition were labelled as “false alarm”. (2) Memory retrieval: trials for which participants correctly recalled the auditory speech stimulus associated with the visual cue were labelled as “hit”. In contrast, trials for which participants did not recall the correct auditory stimulus were labelled as “miss”.

### MEG acquisition

MEG data were recorded using a 306-sensor TRIUX MEGIN system with 102 magnetometers and 204 orthogonal gradiometers in a magnetically shielded room. The data were band-pass filtered online using anti-aliasing filters from 0.1 to 330 Hz and sampled at 1000 Hz. The location of the three fiducial points (i.e., nasion, left and right preauricular points), as well as four head position indicator coils (HPI: behind left and right ears right below hairline, and another two on the forehead with a respective distance of at least 3 cm distance) were digitized using a Polhemus Fastrack electro-magnetic digitizer system (Polhemus Inc., USA). Additionally, approximatively 200 extra points on each participant’s scalp were also digitally sampled to spatially co-register the offline MEG analysis with individual structural MRI image or templates. MEG Data were acquired with participants in a sitting position (MEG gantry at a 60°angle).

### MRI acquisition

Structural MRI images (T1 weighted) were acquired for 17 participants using a 3-tesla Siemens Magnetom Prisma scanner (MP-RAGE, TR = 2000 ms, TE = 2.01 ms, TI = 880 ms, flip angle = 8°, FOV = 256 × 256 × 208 mm, 1 mm isotropic voxel). For the remaining 6 participants without individual T1-weighted structural MRI image, we used the MNI template brain (Montreal, Quebec, Canada) from Fieldtrip.

### MEG preprocessing

Data analysed were performed in Matlab R2018 (MathWorks Inc., USA) by using the Fieldtrip toolbox (Oostenveld et al., 2011) and custom-made scripts. Only gradiometer data were included in the analysis. Data were first bandpass filtered at 0.1-165 Hz to exclude the signal generated by the HPI coils. Second, the continuous data were epoched at the events of interest (encoding: movie onset; retrieval: visual cue onset, first and second auditory speech onsets; Localiser tasks: silent movie and auditory speech onsets), with each epoch beginning 2000 ms before stimulus onset and ending 2000 ms after stimulus onset (total epoch duration: 9000 ms). The duration of the epochs was longer than the windows of interest to exclude potential filter artefacts but were subsequently restricted to windows of interest in the centre of the epochs when conducting statistical analysis. Third, eye-blink or cardiac components were identified by applying independent components analysis and removed. Finally, data were visually inspected, and any artefactual epochs or sensors were removed from the dataset (mean percentage of trials removed: 10%; range: 1-25%; mean number of malfunctioning sensors removed: 1.43; range: 0-6).

### Movement correction

Participants with extreme head motion during MEG recordings were identified as follows: (1) Participants’ data were high-pass filtered to 250Hz to isolate the cHPI signal. (2) For each sensor, the variance of the signal was calculated across time points of the continuous data. (3) The mean variance was averaged across sensors to provide the individual estimate of change in cHPI signal. (4) The mean variance and its standard deviation were calculated across participants. (5) Data sets with a variance greater than three standard deviations above the group mean were excluded from analysis.

### Data reconstruction in the source space

Preprocessed data were reconstructed in the source space using individual head models and structural MRI (T1-weighted) scans for 17 participants. For the 6 participants without structural MRI scans, we used a standard head model and MRI scan templates from the Fieldtrip toolbox. Data was reconstructed using a single-shell forward model and a Linearly Constrained Minimum Variance beamformer (LCMV; van Veen et al., 1997). The lambda regularisation parameter was set to 1%. The source model covered the whole brain with virtual electrodes spaced 10mm apart in all planes.

### Time-frequency decomposition and statistics

Time-frequency decomposition was performed at source level on the encoding and cueing epochs as follows: (1) The preprocessed data was convolved with a 6-cycle wavelet (from −1 s to +5 s with respect to the stimulus onset, in steps of 50 ms; 1-40 Hz; in steps of 1 Hz). (2) The background 1/*f* characteristic was subtracted using an iterative linear fitting procedure to isolate oscillatory contributions (Griffiths et al., 2021; Manning et al., 2009; Zhang and Jacobs, 2015). A vector containing values of each derived frequency *A* and another vector containing the power spectrum averaged over all time-points and trials of the encoding *B*, were log-transformed to approximate a linear function. The linear equation *Ax* = *B* was solved with least-squares regression, in which *x* is an unknown constant describing the 1/*f* characteristic. The 1/*f* fit (*Ax*) was then subtracted from the log-transformed power spectrum (*B*). An iterative algorithm removed peaks in this 1/*f*-subtracted power spectrum that exceeded a threshold determined by the mean value of all frequencies sitting below the linear fit. As the power spectrum is the summation of the 1/*f* characteristic and oscillatory activity, every value below the linear fit can be seen an estimate error of the fit. Any peaks exceeding this threshold were removed from the general linear model, and the fitting was repeated. This step was applied to avoid fit bias due to outlying peaks (Haller et al., 2018).

### Difference of theta power in the whole brain during movie encoding

We limited the analysis to a time-window of +1 s to +4 s with respect to the movie onset to avoid biases induced by the onset and offset of the movie presentation. For every trial, the power spectrum was centred on the frequency corresponding to the peak of MI between lip and auditory envelope signals determined in the movie analyses (± 3 Hz). This step was done to be able to average all the trials together considering the main theta activity carried in each individual movie. For instance, if lip and envelope aligned best at 4 Hz in a movie *n*, the realigned spectrum of the MEG epoch corresponding to the encoding of movie *n* was now 4 ± 3 Hz (from 1 to 7 Hz; 1 Hz bin), ensuring that the central bin of each single trial corresponds to the objectively determined frequency peak of theta activity. Then, the realigned theta power spectrum was averaged across trials within every participant for group analyses. For each participant, we calculated the difference of theta power between the synchronous minus asynchronous condition (theta_sync_ - theta_async_) in the central bin of the realigned spectrum at all virtual sensors. The difference of theta power at group level was statistically assessed against zero by performing a one-tailed cluster-based permutation test (2000 permutations, alpha threshold = 0.05; cluster alpha threshold = 0.05; minimum neighbourhood size = 3). Cohen’s *d_z_* was used as the measure of effect size for significant clusters only as follows: *d_z_* = mean t-statistic within the cluster divided by the square root of the number of participants (Lakens, 2013). The anatomical regions with maximum voxel activation were defined using the automated anatomical labelling atlas (AAL; Tzourio-Mazoyer et al., 2002).

### Difference in theta power between synchronous and asynchronous stimuli in the hippocampus during encoding

For every participant, the mean difference of theta power theta_sync_ - theta_async_ was averaged across the hippocampal sources. Hippocampal virtual sensors (left + right) were defined using the *wfupickatlas* toolbox for SPM (hippocampal region of interest = 32 virtual sensors). The mean difference of theta power across participants was statistically assessed against the null hypothesis (t = 0) by applying a one-sample t-test (one-tailed). To further address whether the difference of theta power was specific to the dominant frequency aligning lip movements and auditory envelope in the stimuli, we compared the original hippocampal difference of theta power against the hippocampal differences in power of neighbouring frequencies as follows: (1) The power was extracted in the frequencies corresponding to the ones determined with the mutual information analyses, divided or multiplied by 2 times the golden mean 1.618 to generate low (4/3.24 = 1.24 Hz; 5/3.24 = 1.55 Hz; 6/3.24 = 1.85 Hz; 7/3.24 = 2.16 Hz; 8/3.24 = 2.47 Hz) and high (4x3.24 = 12.94 Hz; 5x3.24 = 16.18 Hz; 6x3.24 = 19.42 Hz; 7x3.24 = 22.65 Hz; 8x3.24 = 25.88 Hz) neighbouring frequencies. Dividing or multiplying by two times the golden mean ensured that the low-high neighbour frequencies did not share any harmonic with the original frequencies (Pletzer, Kerschbaum and Klimesch, 2010). (2) As for the original data, the power spectrum for each trial was re-centred on the corresponding low or high frequency to create two temporary data, i.e., power in the low or high frequency band. The realigned power was averaged across trials in the synchronous and asynchronous conditions separately, across hippocampal virtual sensors in the low [1.24-2.47 Hz] and high [12.94-25.88 Hz] bands. The difference of power between synchronous minus asynchronous conditions was computed and averaged across low and high neighbouring frequencies for every participant. Finally, the difference of power difference between the original theta and the neighbouring frequencies was statistically assessed by means of a one-sample t-test (one-tailed).

### Replay of auditory speech envelope during retrieval using theta phase similarity

To test whether theta phase specific patterns reflecting auditory speech envelope are replayed during subsequent retrieval (i.e., cued by visual stimuli) we calculated the similarity of theta phase of the speech envelope of the physical stimulus and the theta phase at retrieval in the auditory cortex. Because it was not possible to determine exactly when participants retrieved speech information during successful trials, we adopted a sliding window approach that allows to detect dynamic memory replay with different onset times (Michelmann, Bowman & Hanslmayr, 2016). This method quantifies the phase similarity for a frequency of interest between the pairs of neural (MEG) and auditory envelope time-series with a sliding window as follows: (1) For each trial, the phase values of every time-point from the two time-series were extracted by multiplying the Fourier-transformed data with a complex Morlet wavelet of six cycles, and down-sampled to 64 Hz (from −1 s to +5 s with respect to the onset). Importantly here, the theta frequency of interest for each trial corresponded to the frequency aligning best lip movements and auditory envelope in the corresponding stimulus. (2) For the first time-point of the MEG time-series (*t1_meg_*), the theta phase similarity was computed between the 1-second window centred on *t1_meg_* (± 500 ms) and a fixed 1-second window centred on the first time-point of the auditory speech envelope time-series corresponding to the same speaker (*t1_env_* ± 500 ms). Phase similarity between *t*1_meg_ and *t1_env_* windows was assessed following the single-trial phase locking value approach (Lachaux et al., 2000; Mormann et al., 2000). To do so, the cosine of the absolute angular distance was computed for each time-point, and the similarity was quantified as 1 minus the circular variance of phase differences over time within the *t1_meg/env_* window. Therefore, a unique phase similarity value between 0 and 1 was obtained for *t1_meg_*. The exact same operation (2) was repeated between the fixed *t1_env_* window and a same-length window shifted by one time-point over the MEG time-series, corresponding to a 1-second window centred on *t2_meg_* (± 500 ms). At the end of the sliding window process, we obtained the phase similarity between all time-points of the MEG epoch (from −1 s to +5 s with respect to the stimulus onset), and the first time-window *t1_env_* of the auditory envelope signal. The exact same operation was then repeated for the remaining time-windows of the envelope signal to generate a matrix of theta phase similarity in the two dimensions (i.e., MEG and auditory envelope signals), at every virtual sensor contained in the two regions of interest (i.e. left and right auditory cortex). The virtual sensors contained in the left and right primary auditory cortices of interest were defined by using the *wfupickatlas* toolbox for SPM (left Heschl’s gyrus = 12 virtual sensors; right Heschl’s gyrus = 12 virtual sensors).

In a first step, we applied this method to establish whether replay of speech stimuli took place during retrieval, independently from the condition (synchronous vs asynchronous) of speech encoding. To address this question, we tested phase similarity during visual cueing against chance level as follows: (1) For every participant, the original theta phase similarity was computed from synchronous and asynchronous trials together (the number of trials from the two conditions was counterbalanced by taking the smallest number of available trials between synchronous and asynchronous). (2) The original theta phase similarity matrix was averaged across trials and across virtual sensors from the Heschl’s gyrus (left and right together). (3) In parallel, one hundred permuted data were generated by applying steps (1) and (2) after mismatching the corresponding pairs of neural and stimulus signals. For every trial, the theta phase similarity was computed between the corresponding MEG signal and a random auditory envelope signal. The auditory envelope was randomly selected from the pool of stimuli with the same dominant theta frequency (i.e., theta frequency aligning best lip movements and auditory envelope), excepting the one corresponding to the same MEG epoch. The permuted theta phase similarity matrix was averaged across trials and sources contained in the Heschl’s gyrus (left + right). This step was repeated one hundred times to generate 100 permuted phase similarity data per participant. (4) The permuted theta phase similarity matrix was averaged across the 100 permuted phase similarity data to obtain a phase similarity value per participant under the null-hypothesis. (5) The difference between the original and the permuted theta phase similarity matrices was statistically assessed at group level with a one-tailed cluster-based permutation test (time-window: from −0.5 s to +5 s with respect to the cueing onset; 2000 permutations, alpha threshold = 0.05; cluster alpha threshold = 0.05).

In a second step, we then tested whether the replay of auditory speech was modulated by audiovisual synchrony during encoding as follows: (1) Theta phase similarity matrix was averaged across trials in the synchronous and asynchronous conditions separately, and across virtual sensors (left and right Heschl’s gyrus separately) for every participant. The number of trials averaged within condition was counterbalanced with a subsample corresponding to the smallest number of available trials between synchronous and asynchronous conditions (mean trial number in synchronous condition: 33.09 ± 6.10 and asynchronous condition: 31.17 ± 6.58). Finally, the theta phase similarity difference between synchronous and asynchronous retrieval (from 0 s to +5 s with respect to the cueing onset) was statistically assessed at group level by means of a cluster-based permutation test (2000 permutations, alpha threshold = 0.05; cluster alpha threshold = 0.05; one-tailed). To source localise the theta phase similarity difference during the successful retrieval, the phase similarity was averaged across every time-point contained within a time-window centred on the maximum t-value coordinates from the significant cluster revealed in the left auditory cortex (± 500 ms in the MEG and auditory envelope dimensions), for the synchronous and asynchronous conditions separately. The difference of theta phase similarity between synchronous and asynchronous retrieval was then statistically assessed by performing a one-tailed cluster-based permutation test at group level (2000 permutations, alpha threshold = 0.05; cluster alpha threshold = 0.05).

### Theta phase similarity between MEG and auditory speech envelope during movie perception

To established where cortical neural oscillations tracked theta activity carried by the auditory speech envelope during the encoding of the movies, we applied the exact same method as described above but for the encoding phase, i.e. when participants perceived the movies. We computed the phase similarity between the MEG signal and the corresponding envelope signal of the epoch within the central time-window, i.e., from +1 s to +4 s with respect to the movie onset). For every trial, the phase similarity was also computed between the MEG signal and a different auditory envelope signal randomly selected from the pool of stimuli with the same dominant theta frequency, excluding the one corresponding to the same MEG epoch. The mean phase similarity was then averaged across every time-point of the central time-window in the matching and unmatching cases of the synchronous and asynchronous conditions separately. The difference of averaged phase similarity across the whole brain between the matching and unmatching signals was assessed by means of a one-tailed cluster-based permutation test for the synchronous condition first, then for the asynchronous condition (2000 permutations, alpha threshold = 0.05; cluster alpha threshold = 0.05; averaged across virtual sensors).

## Supplemental Information

**Figure S1.**
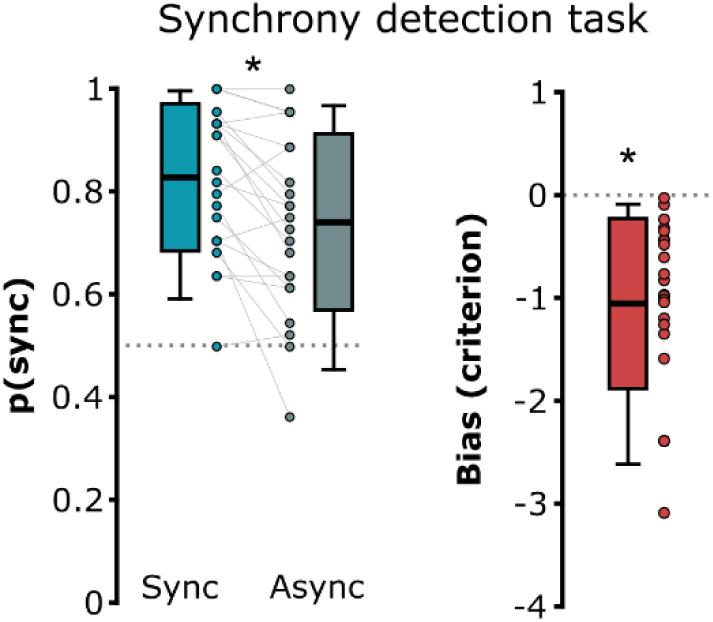
Participant performances in the audio-visual synchrony detection task. (Left) Probability to answer “synchronous” in the synchronous and asynchronous conditions *p*(sync). Participants had a high probability to respond “synchronous” independently from the condition, suggesting that they were actually guessing. The chance level at 0.5 is depicted with the grey dashed line. (Right) The negative bias confirmed that participants tended to respond more systematically “synchronous” independently from the actual (a)synchrony in the movies. The boxplots represent the mean ± standard deviation and the errors bars indicate 5th and 95th percentiles. The dots represent individual means. Significant contrasts are evidenced with stars (p < 0.05).

In complement to the *d’* sensitivity (see Figure 1C), we computed the probability of participants to respond “synchronous” during the audio-visual synchrony detection task (**Figure S1**). If they were sensitive to the audio-visual synchrony in the movies, this probability should drop below chance level in the asynchronous condition (i.e., 0.5). Instead, results showed that participants responded “synchronous” above chance level in both cases (mean *p*(sync)_sync_ = 0.828 ± 0.143 and mean *p*(sync)_async_ = 0.741 ± 0.172). Two one-sample t-test against 0.5 confirmed that both *p*(sync) were significantly above chance (Sync: *t*(22) = 11.022; *p* < 0.001; Cohen’s *d_z_* = 8.189; Async: *t*(22) = 6.736; *p* < 0.001; Cohen’s *d_z_* = 2.13; one-tailed). A paired-sample t-test revealed a statistical difference of *p*(sync) between the two conditions (*t*(22) = 3.956; *p* < 0.001; Cohen’s *d_z_* = 0.549; two-tailed). To confirm that participants were indeed biased towards responding “synchronous” and were virtually insensitive to audio-visual synchrony signal, we computed their individual *c* criterion (mean *c* = −1.054 ± 0.827). A one-sample t-test against zero confirmed that the averaged criterion was significantly below zero and that participants adopted a strategy in responding more systematically “synchronous” across trials (*t*(22) = −6.112; *p* < 0.001; Cohen’s *d_z_* = 1.802; two-tailed). Together with their low *d’*, these results strongly suggest that participants were not sensitive to audio-visual asynchrony during movie encoding as expected.

**Figure S2.**
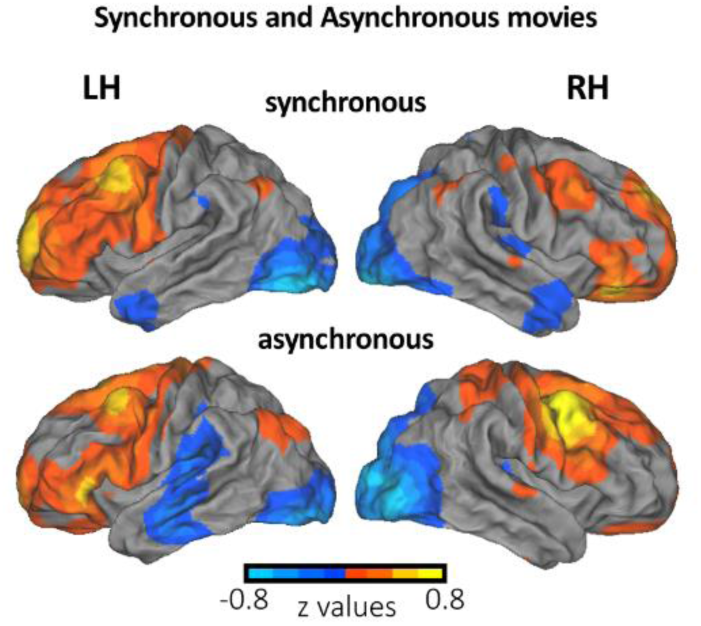
Theta oscillations in the neocortex during synchronous and asynchronous movie encoding. Source localisation of theta power during the encoding of synchronous and asynchronous movies relative to the pre-stimulus baseline (uncorrected z values).

When comparing the theta power responses at the centre of the epoch (i.e., from +2.3 s to +2.8 s with respect to the movie onset) to the pre-stimulus baseline (from −0.7 s to −0.2 s) in the separate synchronous and asynchronous movies, both conditions revealed a predominant increase of bilateral activity in the prefrontal gyrus and in the pre/post central regions. In contrast, the encoding of movies induced also a decrease of theta power in the occipital and the anterior temporal regions.

**Figure S3.**
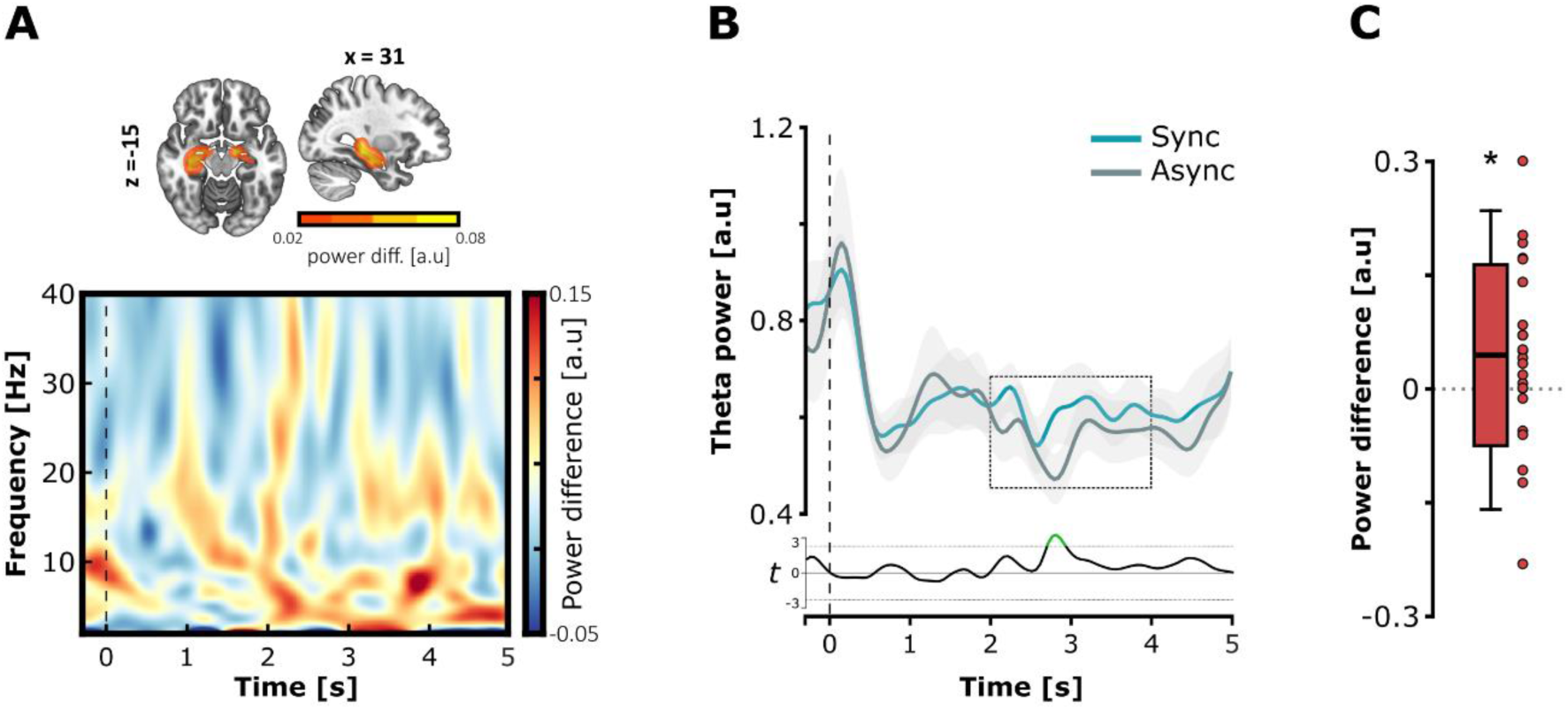
Theta oscillations in the hippocampus during successful retrieval. (**A**) Difference of realigned theta power (*x* = 31; *y* = −19; *z* = −15; top) and time-frequency representation of the difference of spectral power in the hippocampus (bottom) during the visual cueing. The theta oscillations were attenuated when participants recalled correct speech information previously encoded in asynchrony compared to synchrony. (**B**) Temporal modulation of the hippocampal theta power during the retrieval of speech information encoded in the synchronous (blue line: mean ± standard error mean) and asynchronous movies (grey line: mean ± standard error mean). The bottom line (black) depicts the t-values between the two cases for each time-point of the visual cueing (−0.3 s to 5 s with respect to the picture onset). The significant time-points are depicted in green (uncorrected p-values) and served to determine a data-driven time-window of interest for further analyses, i.e., from +2 s to +4 s with respect to the picture onset and centred on the most significant time-point = 2.8 s (dashed rectangle). (**C**) Mean difference of hippocampal theta power in the *post-hoc* time-window (dashed rectangle in B). The successful retrieval of speech information induced a greater theta activity in the hippocampus when audio-visual information was encoded in synchrony as compared to asynchrony in the movies. The boxplots represent the mean ± standard deviation and the errors bars indicate 5th and 95th percentiles. The dots represent individual means. Significant contrasts are evidenced with stars (p < 0.05).

We investigated the theta power responses during the successful retrieval of speech information in the hippocampus beside the phase similarity approach (**Figure S3**). The **Figure S3A (bottom panel)** depicts the time-frequency representation of the difference of power in the hippocampus before realigning the spectrum. Results show that recollecting speech information encoded during asynchronous movies led to a subsequent attenuation of power predominantly localised in the low frequencies. As the exact onset of speech information recollection during visual cueing was not accurately predictable (i.e., varying across trials and participants), we inspected the time course of hippocampal theta power in both conditions to determine when theta responses indexed a difference of memory recollection between speech information encoded in audio-visual synchrony versus asynchrony (**Figure S3B**). The difference was statistically assessed by means of paired-sample t-tests applied at every time-point of the epoch (df = 22; critical value = 1.72; alpha = 0.05; one-tailed). The t-values in function of time revealed a time-window of interest from +2.70 s to +2.95 s with respect to the picture onset, although the p-values were not corrected for multiple comparisons due to the length of the epochs. Nevertheless, we used this information to determine a *post-hoc* time-window of interest centred on the most significant time-point = 2.8 s (*post-hoc* time-window: from +2 s to +4 s with respect to the picture onset) for further analysis. The distribution of mean theta difference between the two conditions in this *post-hoc* time-window was statistically assessed against zero with a one-sample t-test (mean theta power difference = 0.044 ± 0.119). Results revealed that the difference was significantly greater than zero, therefore the retrieval of speech information encoded during synchronous movies induced greater hippocampal theta responses compared to asynchronous movies (*t*(22) = 1.783; *p* = 0.044; Cohen’s *d* = 0.169; one-tailed).

**Figure S4.**
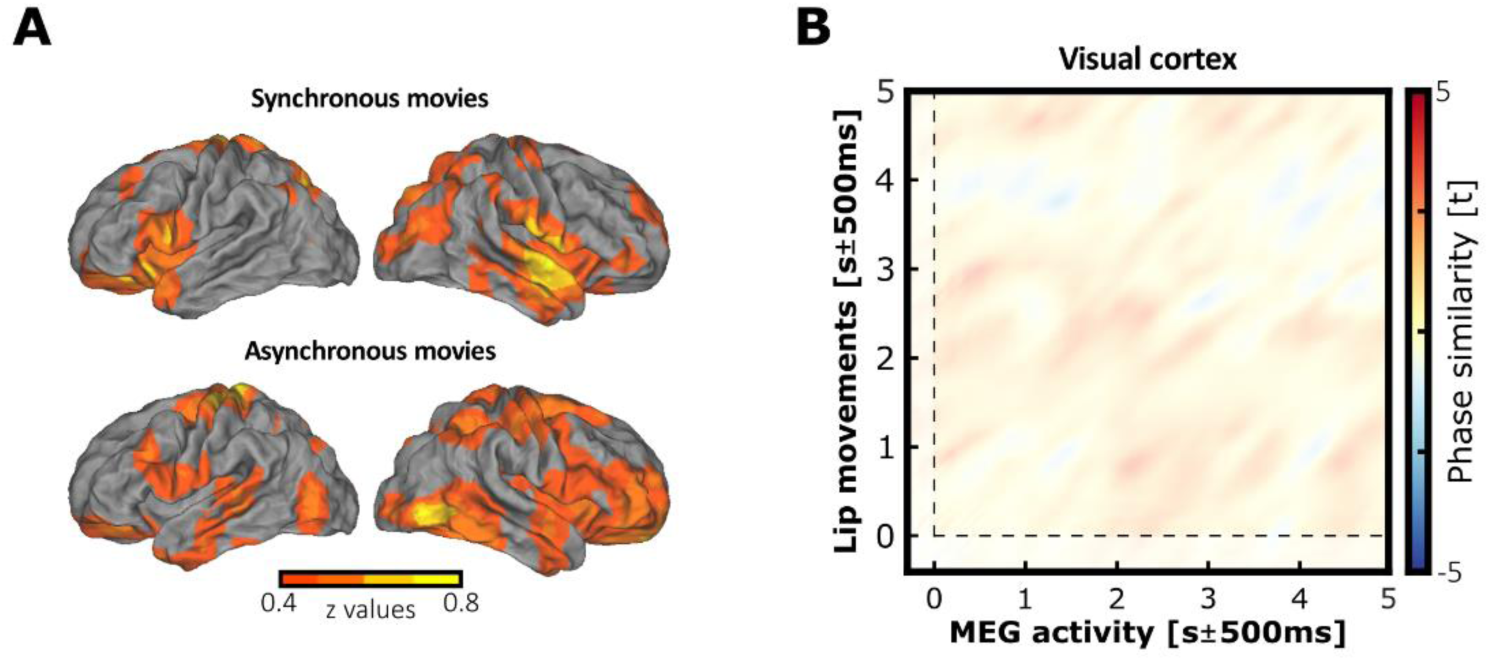
Theta oscillations tracking and reinstatement of the dominant activity carried in the lip movements. (**A**) Source localisation of theta phase similarity between the brain oscillations and lip movements during synchronous or asynchronous movie encoding (threshold at significant t values and normalized for visualisation). In both conditions, the tracking of dominant theta dynamics carried by the lip movements appeared to take place in the expected visual cortex but extended to further regions associated with audio-visual speech integration such as the temporal regions, the inferior-frontal gyrus and the supplementary motor area. (**B**) Theta phase similarity difference between the synchronous versus asynchronous conditions in the visual cortex during successful retrieval. In contrast with the auditory cortex, the fidelity of the theta reinstatement was not significantly modulated by the audio-visual asynchrony in the visual cortex. The non-significant data are masked with transparency.

As for the speech envelope, we analysed how neural theta oscillations tracked the lip movements during movie encoding (**Figure S4A**). We computed the statistical difference of theta phase similarity between corresponding minus random pairs of MEG and lip during synchronous and asynchronous movies. The results revealed a significant cluster when testing the phase similarity difference between corresponding and randomly paired signals in both the synchronous (*p* < 0.001, cluster size = 1.054x10^4^, mean t-statistic within cluster = 3.252; Cohen’s *d_z_* = 0.678) and asynchronous condition (*p* < 0.001, cluster size = 1.258x10^4^, mean t-statistic within cluster = 3.825; Cohen’s *d_z_* = 0.796). No significant negative cluster was found. The theta oscillations tracked lip movements across expected wide-spread networks, including sensory visual cortex, as well as temporal areas. Similar to Figure 4C, we compared whether the theta dynamics conveyed by lip movements were reinstated differently in the visual cortex when participants successfully recalled speech information encoded in synchronous or asynchronous movies. The theta phase similarity between every time-point of the MEG and corresponding lip movement signals was computed in the visual cortex during the correct retrieval (**Figure S4B**). The virtual sensors contained in the visual cortex (Broadmann area 17+18) were defined by using the *wfupickatlas* toolbox for SPM (visual region of interest = 140 virtual sensors). Results revealed no significant difference between synchronous and asynchronous retrieval in the visual cortex, suggesting that the fidelity of lip movement reinstatement by the visual cortex was not notably affected by the theta audio-visual asynchrony during the movie encoding.

**Figure S5.**
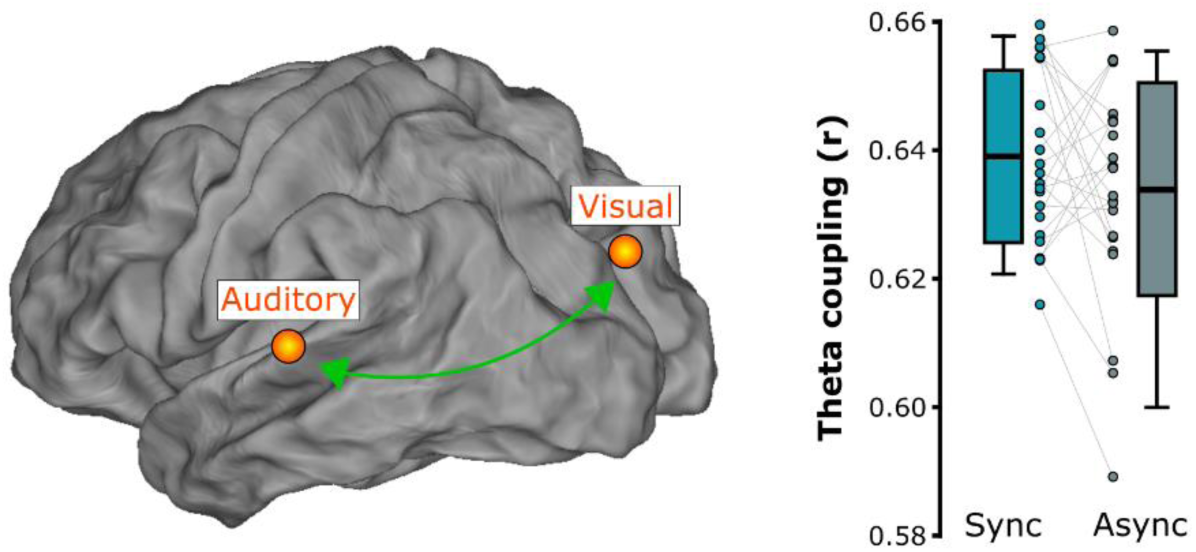
Theta phase coupling between the visual and left auditory cortices during movie encoding. (Left) Visualisation of the sources of interest located in the left auditory cortex and in the visual cortex (orange dots). The phase coupling between the theta activities from the two sources assessed the cross-region connectivity strength during synchronous and asynchronous movie presentation. (Right) Mean theta phase coupling between the left auditory and visual regions in the synchronous and asynchronous movies. The boxplots represent the mean ± standard deviation and the errors bars indicate 5th and 95th percentiles. The dots represent individual means.

Although audiovisual speech organises upon theta oscillations, lip and sound signals are far from stationary. Nevertheless, we aimed to test whether theta activities from the left auditory cortex and the visual cortex synchronised during the encoding of audiovisual movies (**Figure S5 Left**). If so, we would expect a greater phase oscillation coupling between the sensory regions in the case of synchronous audiovisual speech, reflecting more optimal cross-region communication driven by the theta timing compared to asynchronous speech (**Figure S5 Right**). First, the auditory source of interest was determined as the source exhibiting the maximal theta phase similarity (t-value) between the MEG signal from the virtual sensors contained in the left auditory cortex, and the auditory envelop during synchronous movie encoding. Similarly, the visual source of interest was determined as the source exhibiting the maximal theta phase similarity (t-value) between the MEG signal from the virtual sensors contained in the visual cortex and the lip movements during synchronous movie encoding. Second, the signal from left auditory source (*S3585*) was projected orthogonally onto the visual signal (*S4223*) applying a Gram-Schmidt process for single trials before computing phase information. This was done to reduce the noise correlation patterns reflecting activity from a common source estimate captured at different electrodes.

Third, for each trial the instantaneous theta phase of the auditory and visual orthogonalized time-series were computed by applying a Hilbert transform with a bandpass filter centred on the dominant frequency bin ± 2 Hz determined individually for every movie. Third, the absolute difference of unwrapped instantaneous phase between auditory and visual sources was computed for each single trial at each time-point in the central time-window, i.e., from +1 s to +4 s with respect to the movie onset. To assess that the theta phase coupling (ϕA-V) between visual and auditory cortices was affected by the theta asynchrony in the movies, we computed and compared the resultant vector length *r* of the distance between the observed ϕA-V in the data and a theoretical angle ϕA-V = 0 in the synchronous and asynchronous conditions separately. For each trial, we calculated *r* from the absolute distance between the real auditory-visual phase offset and the arbitrary phase offset of 0, at each time point of the central time-window for the two separate conditions. The resultant vector length was collapsed across time in the synchronous and asynchronous conditions separately, resulting in two values per trial. Single-trial values in the two conditions were then averaged across trials for every participant, and the absolute difference of phase coupling between synchronous and asynchronous movies was assessed with a paired samples t-test (one-tailed). At first glance, the difference of theta coupling between the left auditory and visual cortices seemed to support our hypothesis (mean vector length *r* Synchronous = 0.639 ± 0.0134; mean vector length *r* Asynchronous = 0.633 ± 0.0167). However, the paired-sample t-test revealed no statistical difference of vector length *r* between the two conditions (*t*(22) = 1.293; *p* = 0.105; Cohen’s *d_z_* = 0.017; one-tailed). Phase-coupling analysis did not establish that disorganising theta timing between speech features was reflected in the across-region communication during movie encoding.

**Figure S6.**
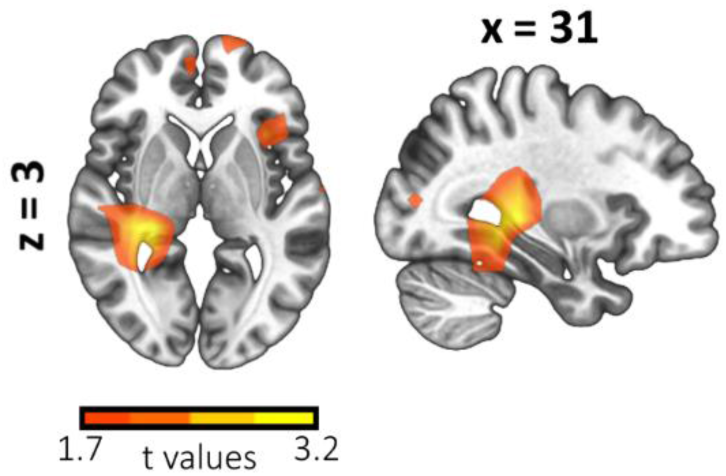
Source localisation of the theta phase similarity difference between the synchronous versus asynchronous conditions during successful movie retrieval. The time-window corresponds to the maximum t-value coordinates ± 500 ms in the MEG activity and speech envelope dimensions of the significant cluster in the left auditory cortex (threshold at t values of the first non-significant cluster).

The **Figure S6** represents the difference in theta phase similarity between the synchronous and asynchronous condition across the whole brain at the maximum t-value coordinates from the above significant cluster in the left auditory cortex (respectively 2.891 s ± 500 ms in the MEG dimension and 1.234 s ± 500 ms in the envelope dimension with respect to signal onsets). Results are only reported for informativeness purpose as no significant cluster was revealed at whole brain level analysis.

Finally, we aimed to verify that the neocortical oscillations tracked the theta oscillations conveyed by the lip movements and the auditory envelope during speech perception. At the end of the memory task, participants completed two short unimodal tasks presenting either a subsample of 60 silent movie (i.e., visual task) or 60 corresponding auditory speech track (i.e., audio task) stimuli taken from the memory task. The visual task included 10 additional target stimuli during which the video froze for five consecutive frames while the speaker’s mouth was either open or closed. Participants were instructed to carefully pay attention to the speaker’s lip movements and to indicate with a button press every time they detected the video freezing when the speaker’s mouth was open during presentation. The audio task also included 10 additional target stimuli, during which a pure auditory tone (1kHz; 100ms with random onset) was embedded in the auditory speech track. Participants were instructed to carefully listen to the auditory stimuli and to indicate with a button press every time they detected a pure tone during speech presentation. In both unimodal tasks, the order of trials (stimuli + targets) was randomised and presented in 10-stimulus blocks separated by short breaks to allow participants to rest. Target trials were excluded from the analyses. Theta coupling between MEG epochs and corresponding lip or envelope signals was quantified by computing Mutual Information (MI) at the dominant frequency determined with movie analyses (see Methods). Theta entrainment in the whole brain was assessed by comparing the MI at the centre of the epochs during silent movie or auditory speech perception (i.e., from +2.3 s to +2.8 s with respect to the stimulus onset) to the pre-stimulus baseline (from −0.7 s to −0.2 s) at each virtual source (**Figure S7**).

**Figure S7.**
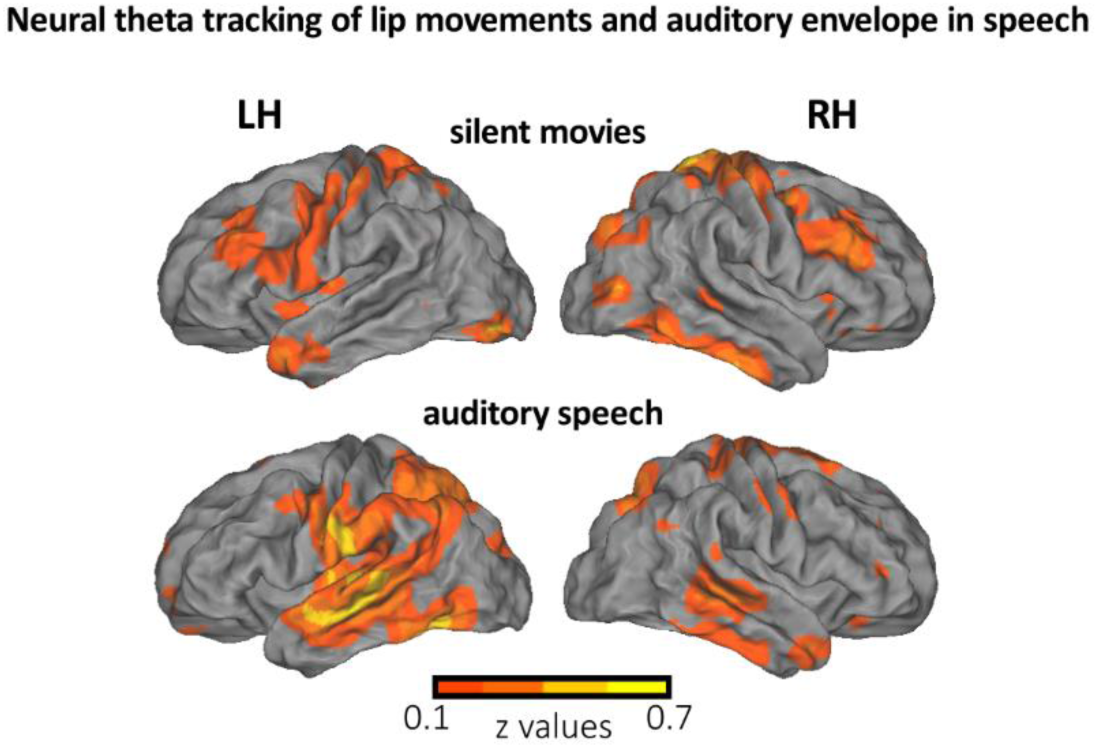
Neocortical activity tracks theta oscillations carried by the lip movements and auditory envelope in speech. Source localisation of Mutual Information (MI) between MEG and lip movements (upper part) or auditory envelope (bottom part) signals during silent visual or auditory speech perception, relative to the pre-stimulus baseline (uncorrected z values).

During silent movie perception (**Figure S7 upper part**), neural theta oscillations synchronised to the silent lip movements in the bilateral occipital regions, bilateral motor regions, paracentral lobule region, as well as the bilateral temporal regions. In contrast, neural theta oscillations preferentially synchronised with the auditory envelope in the temporal regions, including auditory cortices (left dominant), left pre/postcentral and the parietal region during auditory speech perception (**Figure S7, bottom part**).

